# Neuropathic Pain Linked to Defective Dopaminergic Inhibition in Anterior Cingulate Cortex

**DOI:** 10.1101/2020.05.09.086454

**Authors:** Kevin Lançon, Edita Navratilova, Frank Porreca, Philippe Séguéla

**Author notes:** **Correspondence:** Dr. Philippe Séguéla, Montreal Neurological Institute, 3801 University, Suite 778, Montreal, Quebec, Canada H3A 2B4,. **Contributions:** Experiments performed by KL and EN. KL and PS designed the project and all authors wrote the manuscript.

## Abstract

Pyramidal neurons in the anterior cingulate cortex (ACC), a prefrontal region involved in processing the affective components of pain, display hyperexcitability in chronic neuropathic conditions and their silencing abolishes hyperalgesia in rodents. We show here that dopamine, through D1-like receptor signaling, inhibits layer 2/3 pyramidal neurons of mouse ACC. Hyperpolarization-activated cyclic nucleotide-gated (HCN) channels control the firing activity of these pyramidal neurons. Through Gs-coupled D1-like receptors, dopamine induces the opening of HCN channels at physiological membrane potentials, driving a significant decrease in input resistance and excitability. Systemic L-DOPA rescues HCN channel activity, pyramidal excitability in ACC as well as sensory phenotype in neuropathic mice while microinjection of a selective D1-like agonist in ACC induces relief of ongoing pain in neuropathic animals. We conclude that decreased dopaminergic inhibition in ACC plays a critical role in an abnormal top-down modulation leading to neuropathic pain.

## Introduction

Damage to peripheral nerves induces maladaptive changes in both peripheral and central nociceptive pathways that may increase the risk of chronic pain (Campbell et al, 2006; Decosterd et al, 2000). These maladaptive changes can elicit ongoing pain as well as allodynia, a pathological sensitivity to innocuous stimuli, and hyperalgesia, an increased sensitivity to painful stimuli (von Hehn et al, 2012). These hallmark symptoms of chronic pain are linked to an increase in pyramidal cell excitability in layer 2/3 of the anterior cingulate cortex (ACC), a region of the medial prefrontal cortex (mPFC) that is implicated in the affective components of pain in both human and animal studies (Cordeiro Matos et al, 2015; Eto et al, 2011; Kamiński et al, 2017; Mhalla et al, 2010; Rainville et al, 1997). A recent meta-analysis of brain imaging data showed that the ACC is the most consistently activated region in patients suffering from chronic pain consistent with its critical role in higher-order pain processing (Jensen et al, 2016). Additionally, in experimental rodent models of inflammatory and neuropathic pain, optogenetic inhibition of hyperexcitable pyramidal cells has been reported to produce analgesic effects including decreased tactile allodynia and conditioned place preference, respectively (Kang et al, 2015; Sellmeijer et al, 2018). Collectively, these findings support the conclusion that the ACC is a region involved in regulatory top-down pain processing. Despite these findings, very little is known about how pyramidal neurons in the ACC are modulated in physiological or pathological states.

Hyperpolarization-activated cyclic nucleotide gated (HCN) channels control the excitability of many central neurons by regulating input resistance (Poolos et al, 2002). These cAMP-gated cation channels are highly expressed in the mPFC and have decreased open channel probabilities at physiological membrane potentials in chronic pain conditions (Cordeiro Matos et al, 2015; Notomi et al, 2004; Santello et al, 2017). This reduction in HCN-mediated currents (I_h_) results in increased input resistance and therefore excitability (Cordeiro Matos et al, 2015; Santello et al, 2017). Recent findings indicate that HCN channels are colocalized with G_s_-coupled D1-like dopamine (DA) receptors on dendritic spines of mPFC pyramidal neurons (Paspalas et al, 2013; Stoof et al, 1981).

DAergic modulation of the ACC is particularly interesting given the known role of the other major monoamines including norepinephrine and serotonin in prefrontal modulation in chronic pain states (Cordeiro Matos et al, 2015; Santello et al, 2017). In view of the well-documented effects that both norepinephrine and serotonin have on HCN channel function in the ACC and consequently on pathological cortical excitability, we hypothesized that DA, through postsynaptic D1-like receptors and opening of HCN channels, can decrease pyramidal excitability and consequently influence top-down pain pathways by inhibition of pyramidal cells.

In the present study, we investigated the neuromodulatory effects of DA on intrinsic cellular excitability of pyramidal cells in layer 2/3 of the ACC and we provide electrophysiological evidence of a functional interaction between D1-like receptors and HCN channels. We also examined whether an increase in DAergic input to the ACC can reduce the pathological cellular dysfunction of the ACC observed in neuropathic conditions and alleviate the aversive symptoms of chronic pain. We found that a tonic increase in cortical DA, via long-term systemic administration of L-DOPA/carbidopa, relieves neuropathy-induced hypersensitivity while acute intracortical injection of a selective D1-like receptor agonist drives conditioned place preference selectively in neuropathic animals indicating the relief of the aversive state of ongoing pain.

## Results

### DAergic Inhibition of Pyramidal Cells in ACC

Following application of DA (10-50 µM) by perfusion for 15 minutes, 50% (n = 9 of 18) and 54% (n = 7 of 13) of pyramidal cells in layer II/III of the ACC were inhibited relative to baseline (data not shown).The dichotomy in response to DA between responsive and non-responsive cells was obvious, as responsive cells showed a marked inhibition of −40.1 ± 10.1% in contrast to non-responsive cells which expressed a +6.9 ± 4.5% excitation relative to baseline values following 50 µM DA application (Figure 1A) (n = 18, unpaired t-test: t(14)=4.6, p = 0.0004). Criteria for responsiveness was based on the presence of a reversible decrease in excitability. Analysis of responsive cells showed a dose-dependent inhibition. We observed that 10 µM DA reduced excitability to 84.9 ± 6.4% of baseline (15% inhibition) while 50 µM DA inhibited cells to 59.7 ± 8.4% of baseline (40% inhibition, Figure 1B and 1C) (n = 7-9; one-way ANOVA: F(2, 21) = 7.6, p = 0.0032). The inhibitory effects of DA were reversible following a 10-minute washout in all responsive cells, with excitability values for both 10 µM and 50 µM DA treatments returning to within 1.6% of control values (Figure 1C). The input-output plot illustrates the inhibitory effect of DA on pyramidal firing activity (Figure 1C). Interestingly, DA induces an increase in the threshold required to trigger an action potential at low stimulation intensities. This is represented in Figure 1C where weak stimulations below 50 pA no longer elicited action potentials in cells treated with 50 µM DA (n = 7, 9; two-way ANOVA: F(14, 270) = 56.0, p = 0.0001). This is further demonstrated by analyzing the current required to evoke an action potential. We found DA increased the current required to evoke an action potential at −60 mV from ≈40 pA to 56.7 ± 8.8 (10 µM) and 68.9 ± 10.6 pA (50 µM) (n = 9, paired t-test: t(8) = 3.5, p = 0.0101)(Figure 1C). To check whether a decrease in excitability is observed over time in control conditions, we treated pyramidal cells exclusively with aCSF alone and noticed no significant change in excitability (2.0 ± 6.1% relative to baseline values, n=8, see Figure 1C). In concordance with the decreased excitability of layer 2/3 pyramidal cells of the ACC following DA application, we also observed a significant decrease in the input resistance of DA-responsive cells (Figure 1D). Input resistance for cells treated with aCSF alone was found to be 145.7 ± 8.7 MΩ whereas in the presence of 50 µM DA we measured an input resistance of 114.4 ± 10.2 MΩ (Figure 1D) (n = 8, paired t-test: t(7)=2.3, p = 0.028). Similarly, 10 µM DA also induced a trend of reduction in input resistance, however the difference was not found significant (154.8 ± 9.8 MΩ to 128.1 ± 8.7 MΩ) (data not shown). These electrophysiological results indicate that the neuromodulator DA induces a robust and reversible inhibition of pyramidal cells in the mouse ACC, by inducing a decrease in their input resistance.

**Figure 1.**
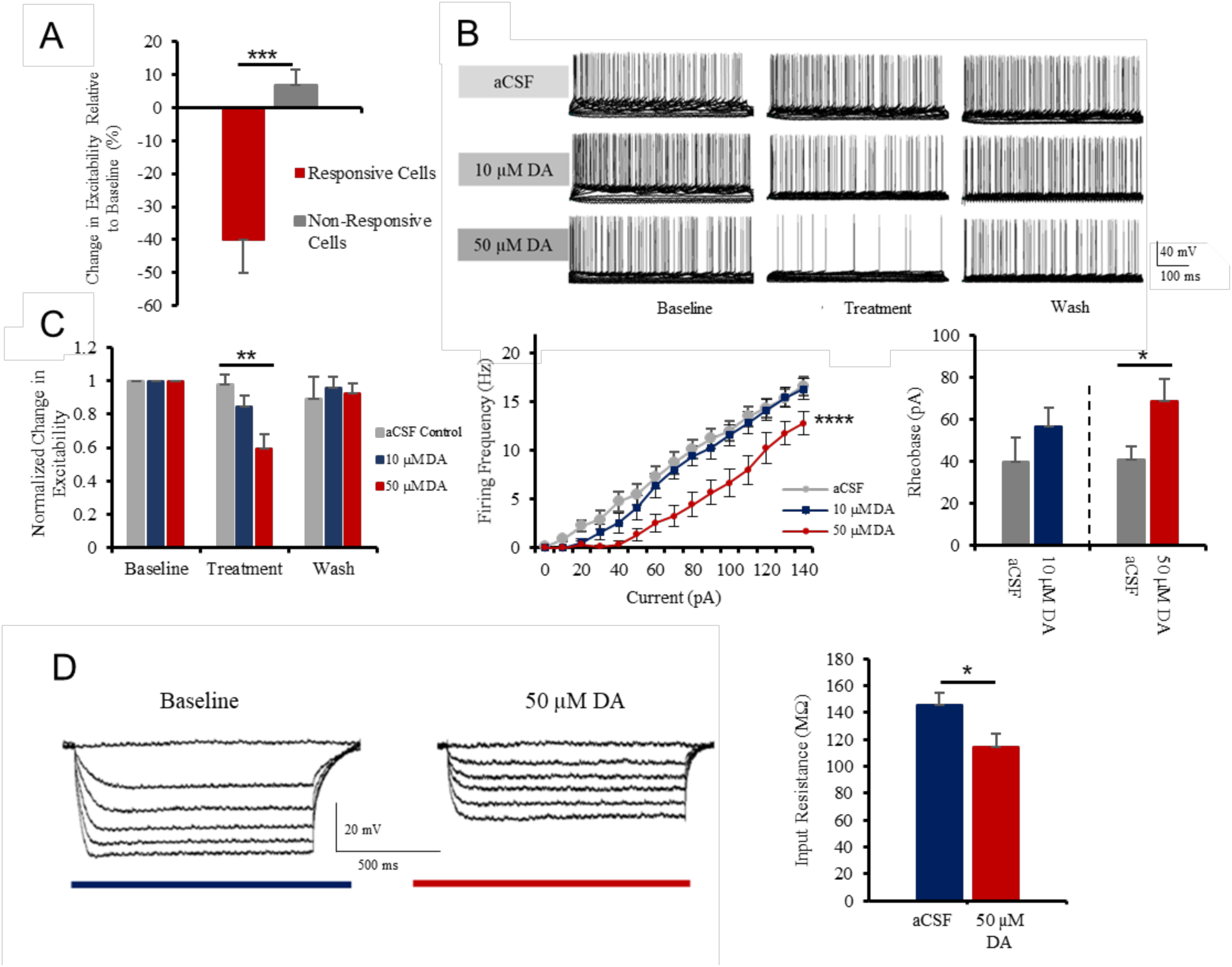
DA inhibits pyramidal cells in layer 2/3 of the mouse ACC. A. Following application of 50 µM DA, a large subset of layer 2/3 pyramidal neurons in the ACC shows strong inhibition. B. 10-50 µM DA reduces excitability in a dose-dependent and reversible fashion, compared to control conditions, electrophysiological current-clamp traces shown. C. Left panel: Quantitative group analysis shows that 50 µM DA significantly inhibits pyramidal firing activity in the ACC in comparison to control conditions. Middle panel: Input-output plot demonstrates interaction between stimulation intensity and firing frequency following treatment with DA. Right panel: Application of 50 µM DA induces a significant increase in pyramidal rheobase. D. Left panel: Typical current-clamp recordings illustrating alterations in input resistance following treatment with 50 µM DA. Right panel: Quantitative group analysis of input resistance of pyramidal cells treated with aCSF or 50 µM DA. Values represented as means ± SEM (n = 9-18): *p < 0.05, **p < 0.01, ***p < 0.001, ****p < 0.0001.

### DAergic Inhibition of Pyramidal Cells Mediated by D1-like Receptors

To determine which receptor DA acts at on pyramidal cells, we tested the effects of the selective D1-like agonist SKF81297 and D2-like agonist quinpirole. Application of 10 µM SKF81297 induces a strong, reversible inhibition of layer 2/3 pyramidal cells in the ACC (Figure 2A), similar to the effects induced by 50 µM DA (Figure 1C). SKF81297 elicited a significant decrease in excitability relative to control values (Figure 2B) (n = 9; control: 98.0 ± 6.1 %, SKF: 68.9 ± 3.2% relative to baseline; one-way ANOVA: F(2, 24) = 16.2, p = 0.0013). SKF81297 also evoked a significant decrease in input resistance (Figures 2C) (n = 9; baseline: 151.2 ± 6.5 MΩ, SKF: 134.7 ± 9.2 MΩ; paired t-test: t(8) = 2.17, p = 0.031).

**Figure 2.**
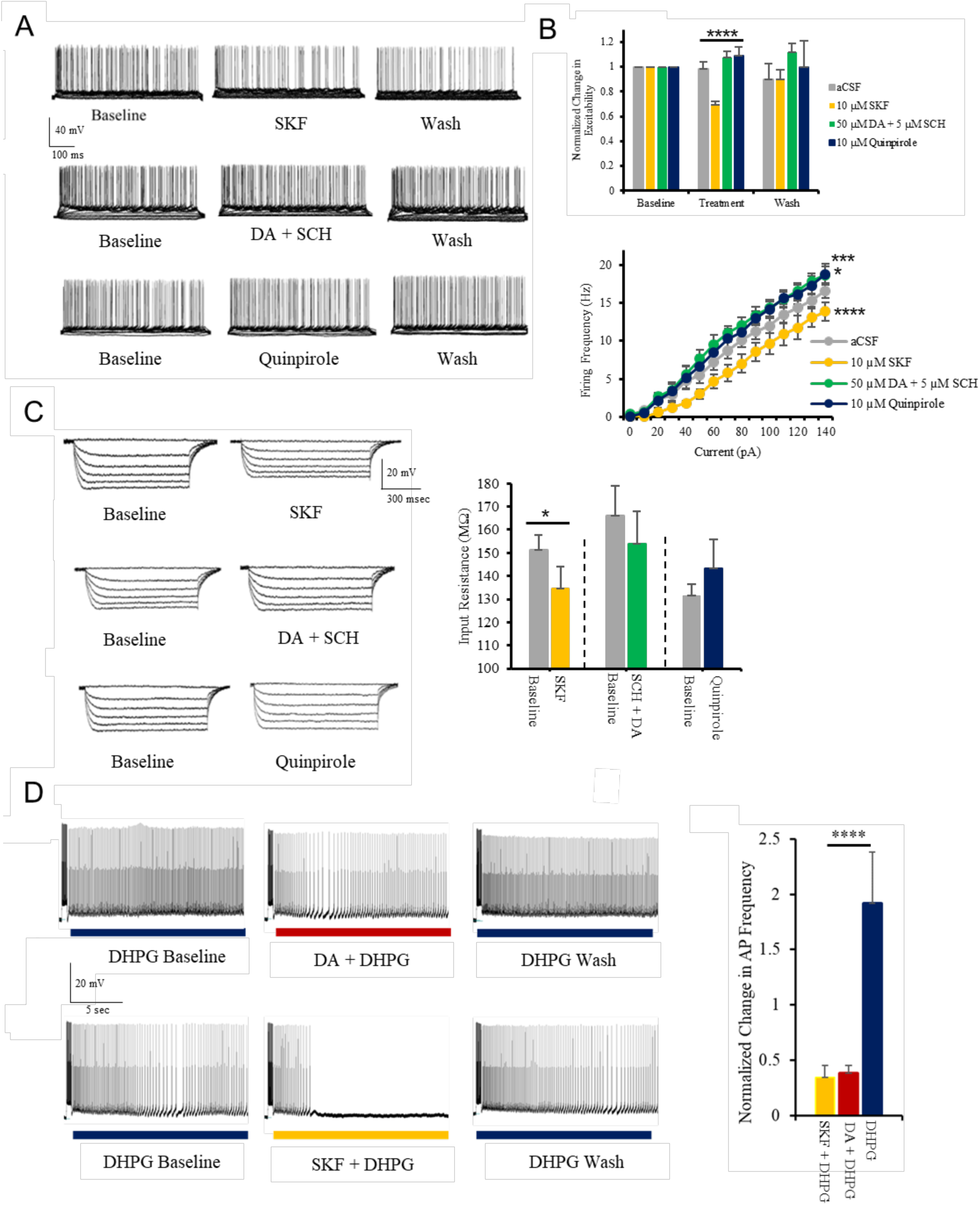
DAergic inhibition of pyramidal cells in ACC is exclusively mediated by D1-like receptors. A. Typical current-clamp recordings illustrating alterations in pyramidal firing activity following application of 10 µM SKF81297, 5 µM SCH23390 + 50 µM DA, and 10 µM quinpirole. B. Top panel: Quantitative group analysis of cell excitability, SKF81297 induces a significant decrease in excitability whereas 5 µM SCH23390 + 50 µM DA and 10 µM quinpirole did not have any inhibitory effect. Bottom panel: Input-output plots recorded during each of 4 different treatments. C. Left panel: Typical traces illustrating the impact of various DAergic ligands on input resistance. Right panel: 10 µM SKF81297 induces a significant decrease in input resistance whereas 5 µM SCH23390 + 50 µM DA and 10 µM quinpirole did not. D. Left panels: Electrophysiological current-clamp traces illustrating 10 µM DHPG-induced persistent firing along with the reversible inhibitory effects of 50 µM DA and 10 µM SKF81297. Right panel: Quantitative group analysis illustrating the significant decrease in firing frequency during persistent firing following application of 50 µM DA or 10 µM SKF81297 in comparison to the control group (DHPG alone). Values represented as means ± SEM (n= 6-10): *p < 0.05, ***p < 0.001, ****p < 0.0001.

We also investigated the effects of SCH23390, a selective D1-like antagonist, on the inhibitory effects of DA. Addition of SCH23390 by pre-incubation and co-application blocks both the inhibitory effect of 50 µM DA (Figure 2A) and the DA-induced decrease in input resistance (Figure 2C). Group analysis further confirms this antagonism as SCH23390 application completely abolishes the effects of DA on excitability (n = 10, +7.6 ± 4.9 % excitability relative to baseline) and on input resistance (n = 10, 166.0 ± 12.8 MΩ to 154.0 ± 13.9 MΩ following DA application) (Figures 2B and 2C). Interestingly, directly blocking D1-like receptors and adenylate cyclase activation via the antagonist SCH23390 caused an increase in baseline excitability (Figure 2B) (n = 10, p = 0.0004), suggesting the presence of basal dopamine levels in ACC.

We then investigated the effect of activating G_i_-coupled D2-like receptors on pyramidal cell excitability (Tsu et al, 1996). We applied the selective D2-like agonist quinpirole (10 µM) to pyramidal neurons in layer 2/3 of the ACC and observed that D2-like receptor activation increases their firing activity (Figure 2A). Following quinpirole application we found a significant increase in excitability in comparison to aCSF controls (Figure 2B) (n = 6, 8; repeated measure two-way ANOVA with Dunnett’s multiple comparison: F(3, 450) = 30.2, p = 0.0448). Although we noticed a trend to increased input resistance, this effect was not found significant (Figure 2C). Altogether, these findings indicate that the postsynaptic DAergic inhibition of pyramidal cells in the ACC exclusively depends on D1-like receptors and does not involve excitatory D2-like receptors.

### DAergic Modulation of Persistent Firing in ACC

Pyramidal cells in the ACC display persistent firing properties that make them capable of holding a working memory trace (Kamiński et al, 2017). We checked if persistent firing was also susceptible to DAergic inhibition. Persistent firing was induced by coincident depolarization and stimulation with the mGluR1/5 agonist DHPG (10 µM). DHPG alone caused an increase of 91.5 ± 46.4% in persistent firing frequency relative to baseline (n = 11) (Figure 2D). In a large majority of persistent firing cells, application of DA (10 µM) or the D1-like agonist SKF81297 (10 µM) produced a significant decrease in firing frequency or even termination of the prolonged firing (n = 12 of 17 for DA, n = 13 of 18 for SKF81297, Figure 2D). SKF81297 and DA application induced a 61.8 ± 10.3% and 65.4 ± 6.9% reduction in persistent firing intensity, respectively, relative to baseline values (n = 12-13; one-way ANOVA: F(2, 30) = 15.3, p < 0.0001) (Figure 2D).

### HCN Channels Mediate DAergic Modulation of ACC Pyramidal Cells

To determine if HCN channels contribute to DAergic inhibitory mechanisms, we pre- and co-applied the HCN channel blocker ZD7288 alongside DA during recordings. We observed that treatment with 10 µM ZD7288 completely abolishes DAergic inhibition in the ACC (Figure 3). Following DA application in presence of ZD7288, we noticed no decrease in either excitability (Figure 3A) or input resistance (Figure 3A and 3B). Whereas 50 µM DA alone induces a 40.3 ± 8.4% decrease in excitability, co-application with ZD7288 evokes a 26.8 ± 12.4% increase in excitability (n = 8, one-way ANOVA: F(2, 22) = 13.4, p = 0.0002) (Figure 3A). Blocking HCN channels suppresses the impact of DA on input resistance (n = 7) (Figure 3B). These findings are in line with previous reports documenting the effect of ZD7288 on the input resistance of central neurons (Surges et al, 2004). To gauge the importance of HCN in maintaining the homeostatic excitability of pyramidal cells in the ACC, we tested the effect of ZD7288 alone, i.e. in the absence of DA. Application of ZD7288 dramatically increases both pyramidal excitability and input resistance to levels observed in neuropathic conditions (n = 8, un-paired t-tests: t(7) = 3.3, 2.5 and p = 0.0053, 0.0245)(Figures 3C and 7A). We found that application of DA or SKF81297 caused a measurable increase in the resting membrane potentials (RMP) of pyramidal cells (n= 17, 15; paired t-test: t(16) = 6.9 and t(14) = 8.4, p < 0.0001 and p < 0.0001) (Figure 3D). This significant shift in RMP was not present when either D1-like receptors or HCN channels were blocked (n = 9, 7, 7) (Figure 3D). Given the nature of HCN channels, the depolarizing shift in RMP could be due to increasing inward cationic currents. Although these results indicate that HCN channels expression and availability are critical for DAergic inhibition, they do not show a functional connection between DAergic modulation and HCN channel-mediated currents (I_h_).

**Figure 3.**
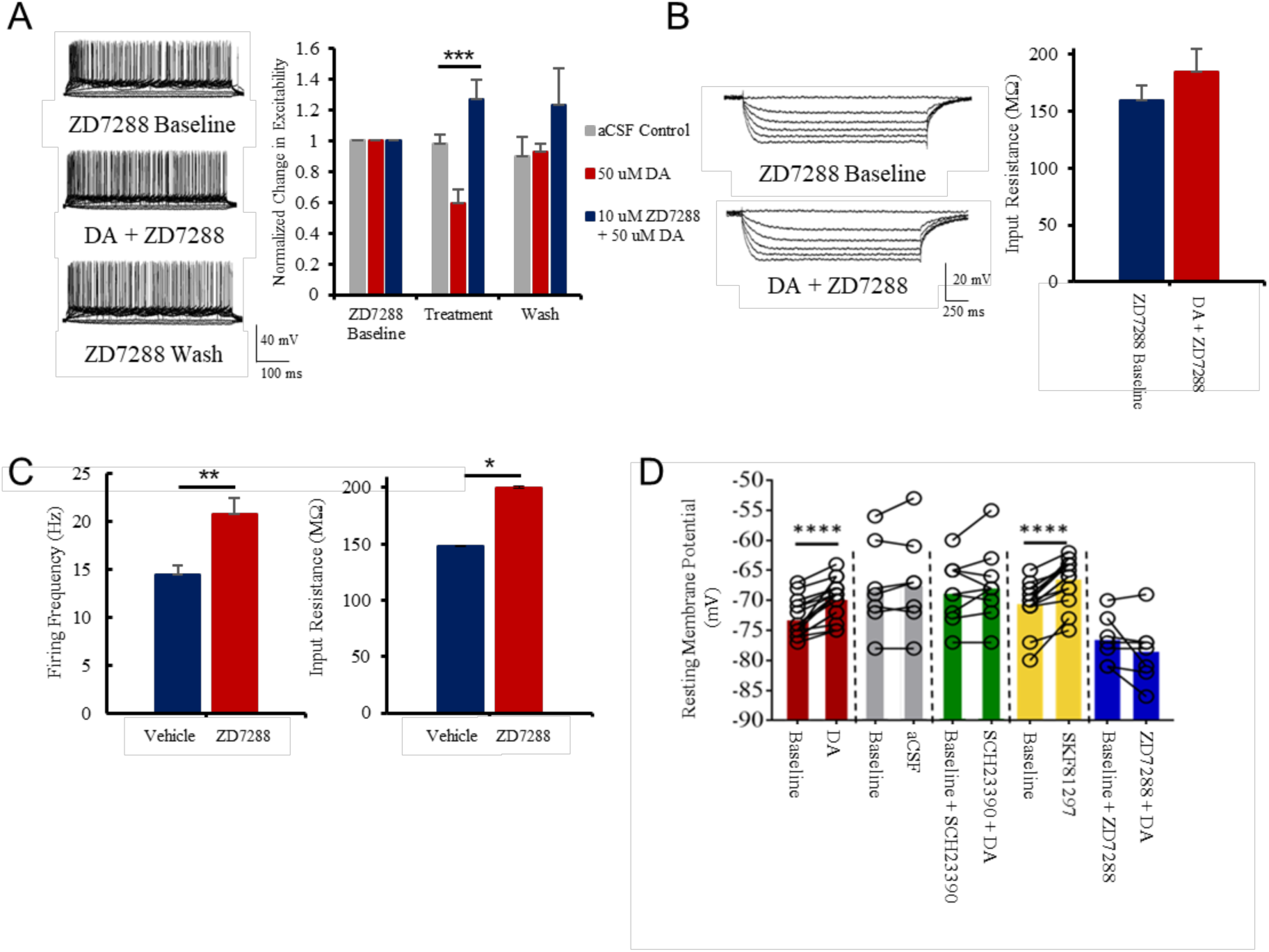
DAergic inhibition depends on HCN channels. A. Treatment with the HCN channel blocker 10 µM ZD7288 occludes the inhibitory effects of 50 µM DA. Representative current-clamp traces showing the effects of 10 µM ZD7288 on excitability (left) and group analysis (right). B. 10 µM ZD7288 blocks the 50 µM DA-induced decrease in input resistance. Representative current-clamp traces showing the effects of 10 µM ZD7288 on input resistance (left) and quantitative group analysis (right). C. Quantitative effects of 10 µM ZD7288 on firing frequency (left) and input resistance (right) at baseline. D. Both 50 µM DA and 10 µM SKF81297 induce a significant increase in resting membrane potentials of layer 2/3 pyramidal neurons of the ACC. This effect is lost in presence of the D1-like antagonist SCH23390 (5 µM) or when HCN channels are blocked with ZD7288 (10 µM). Values represented as means ± SEM (n = 8-17): *p < 0.05, **p < 0.01, ***p < 0.001, ****p < 0.0001.

**Figure 4.**
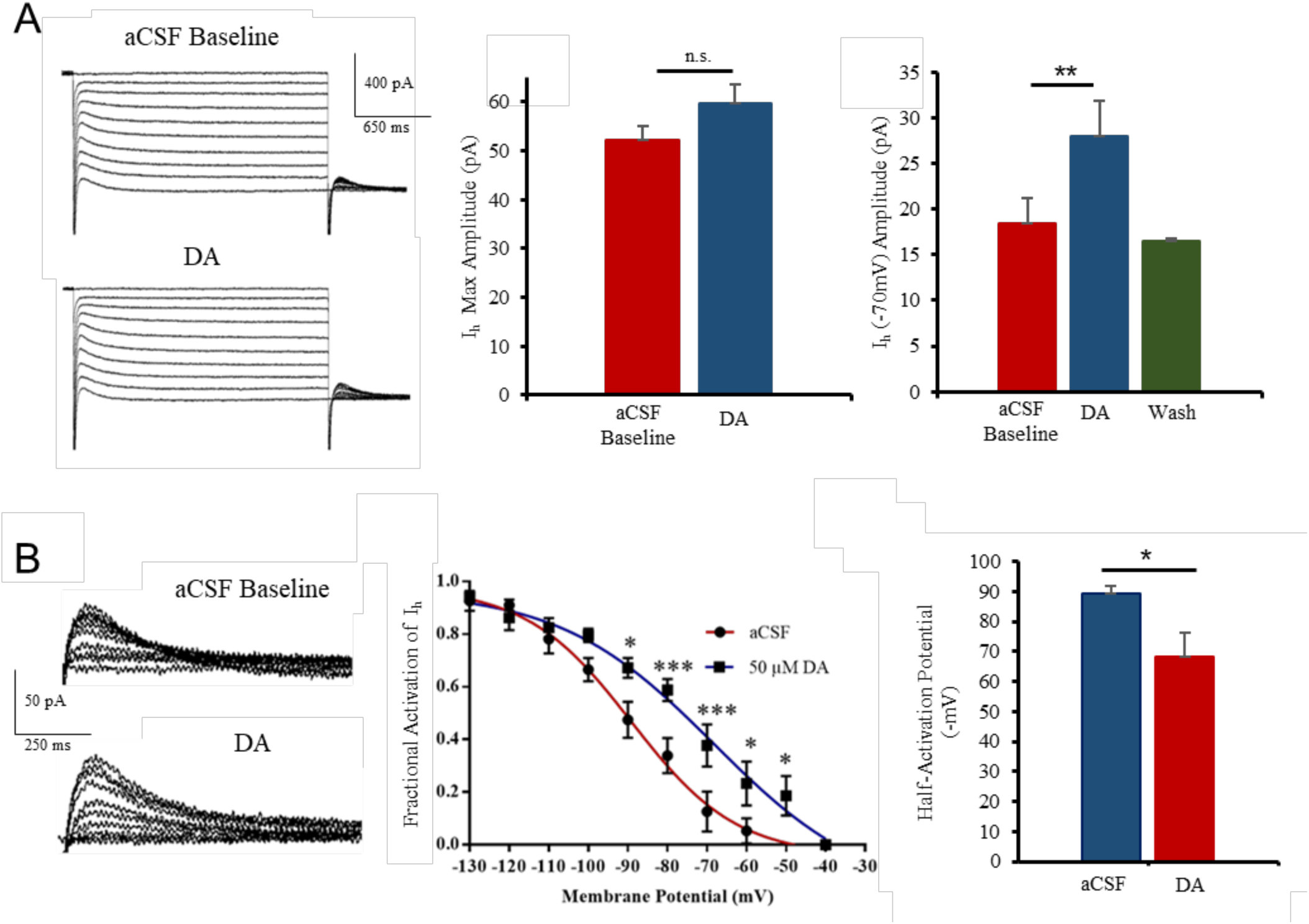
DA controls HCN channel activity in pyramidal neurons of the ACC. A. Left panel: Representative voltage-clamp traces showing HCN currents (holding voltage −130 mV to −40 mV, 10 steps) before and after 50 µM DA application. Middle panel: No significant increase in max I_h_ (recorded at −130 mV) following DA application. Right panel: 50 µM DA induces a strong and reversible increase in I_h_ around physiological membrane potentials (−70 mV). B. Left panel: Typical tail currents shown. Middle panel: Group analysis fitted with Boltzmann function shows a significant increase in open HCN channels at physiological voltages (−90 mV to −50 mV). Right panel: Significant depolarizing shift in half-activation potential of HCN channels after 50 µM DA application. Values represented as means ± SEM (n = 5-9): *p < 0.05, **p < 0.01, ***p < 0.001.

**Figure 5.**
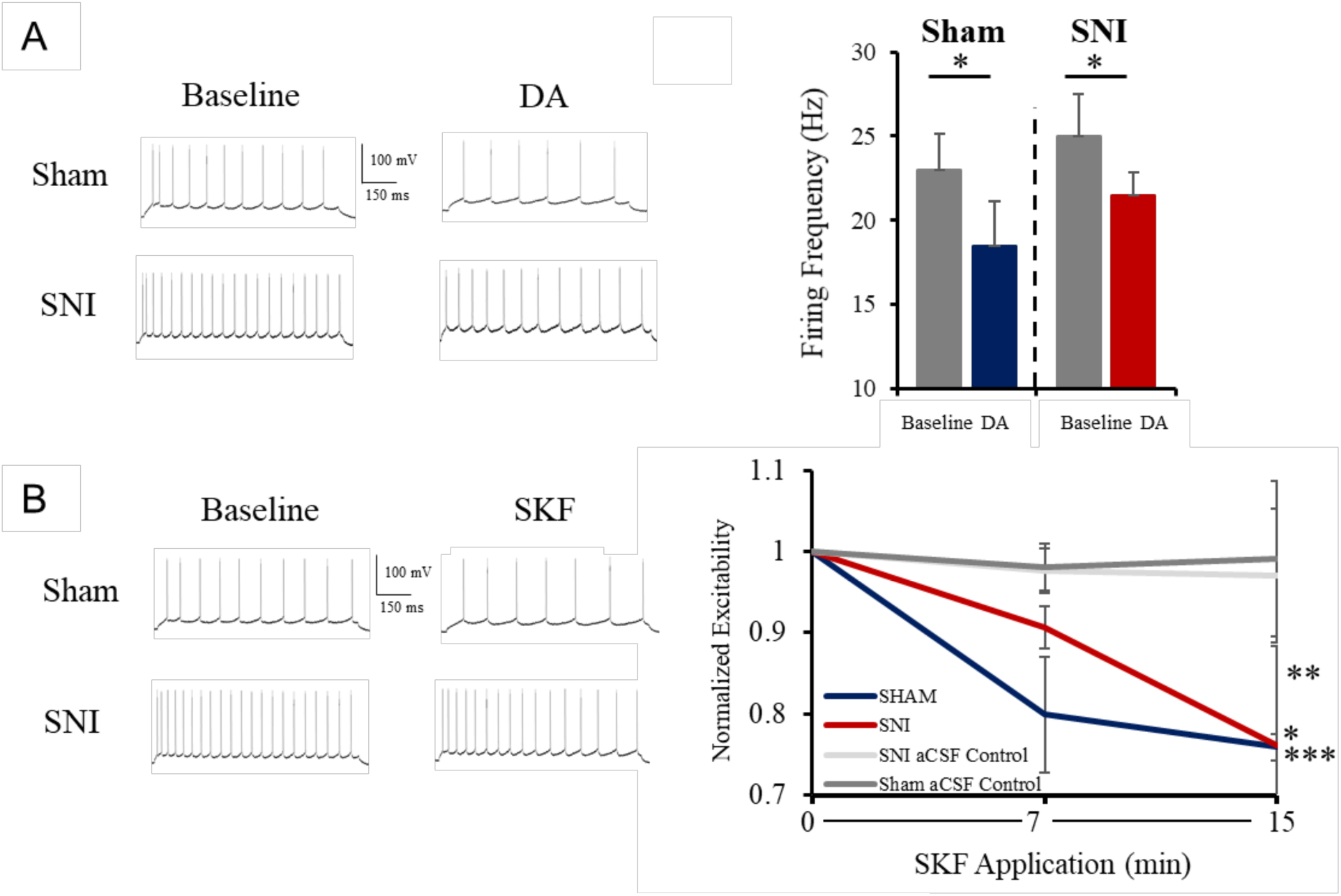
DAergic inhibition is preserved in neuropathic mice. A. Typical current clamp traces illustrating that 50 µM DA induces significant inhibition of layer 2/3 ACC pyramidal neurons in both sham and SNI mice (+40 pA stimulation) (left). At +100 pA stimulation, group analysis shows a significant inhibitory effect of 50 µM DA on firing frequency in sham and SNI mice with similar potency (right). B. Application of 10 µM SKF81297 inhibits layer 2/3 ACC pyramidal neurons in both sham and SNI mice (+40 pA stimulation) (left). Group analysis shows 10 µM SKF81297 significantly decreased excitability relative to baseline and control values (right). Values represented as means ± SEM (n = 4-10): *p < 0.05, **p < 0.01, ***p < 0.001.

**Figure 6.**
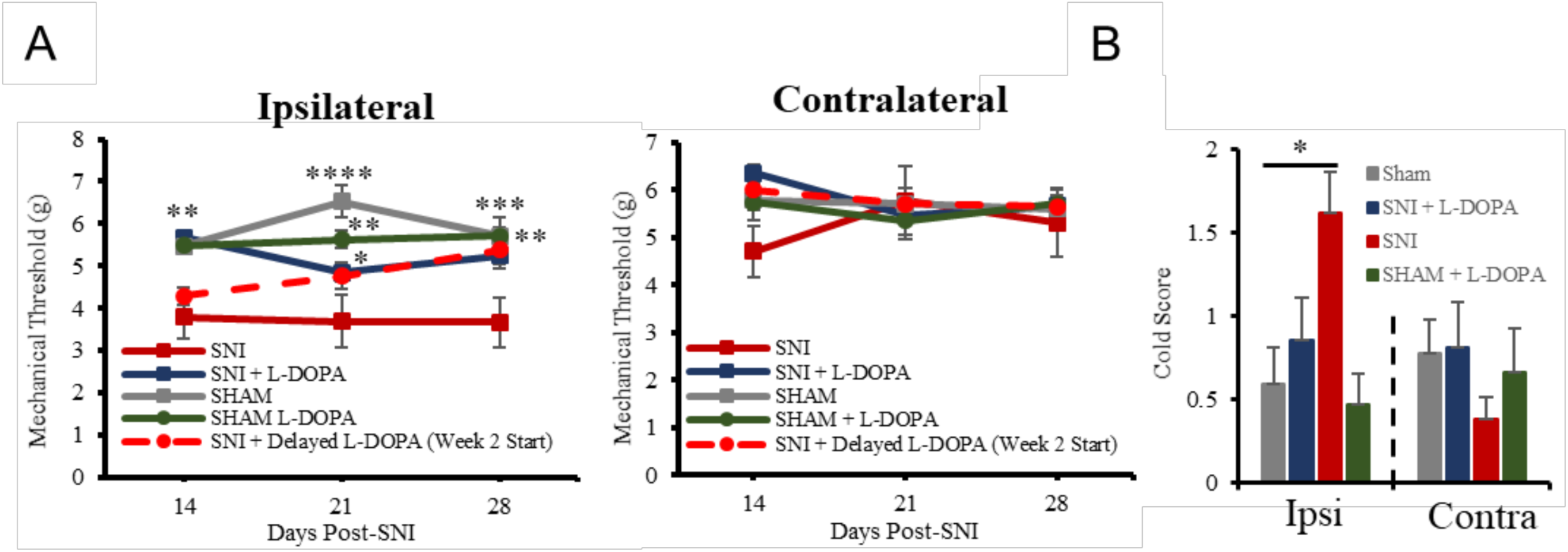
L-DOPA/Carbidopa administration rescues mechanical and cold allodynia in neuropathic mice. A. Mechanical withdrawal thresholds on ipsilateral paw (left) and contralateral paw (right) between treatment conditions. Groups: SNI, sham, SNI + L-DOPA, sham + L-DOPA, and SNI + delayed L-DOPA (supplementation starting two weeks post-SNI surgery rather than on the day of SNI surgery). B. Mean scores of responses to cold in the acetone test on ipsi- and contralateral paws in several treatment conditions. Values represented as means ± SEM (n = 4-10): *p < 0.05, **p < 0.01, ***p < 0.001, ****p < 0.0001.

**Figure 7.**
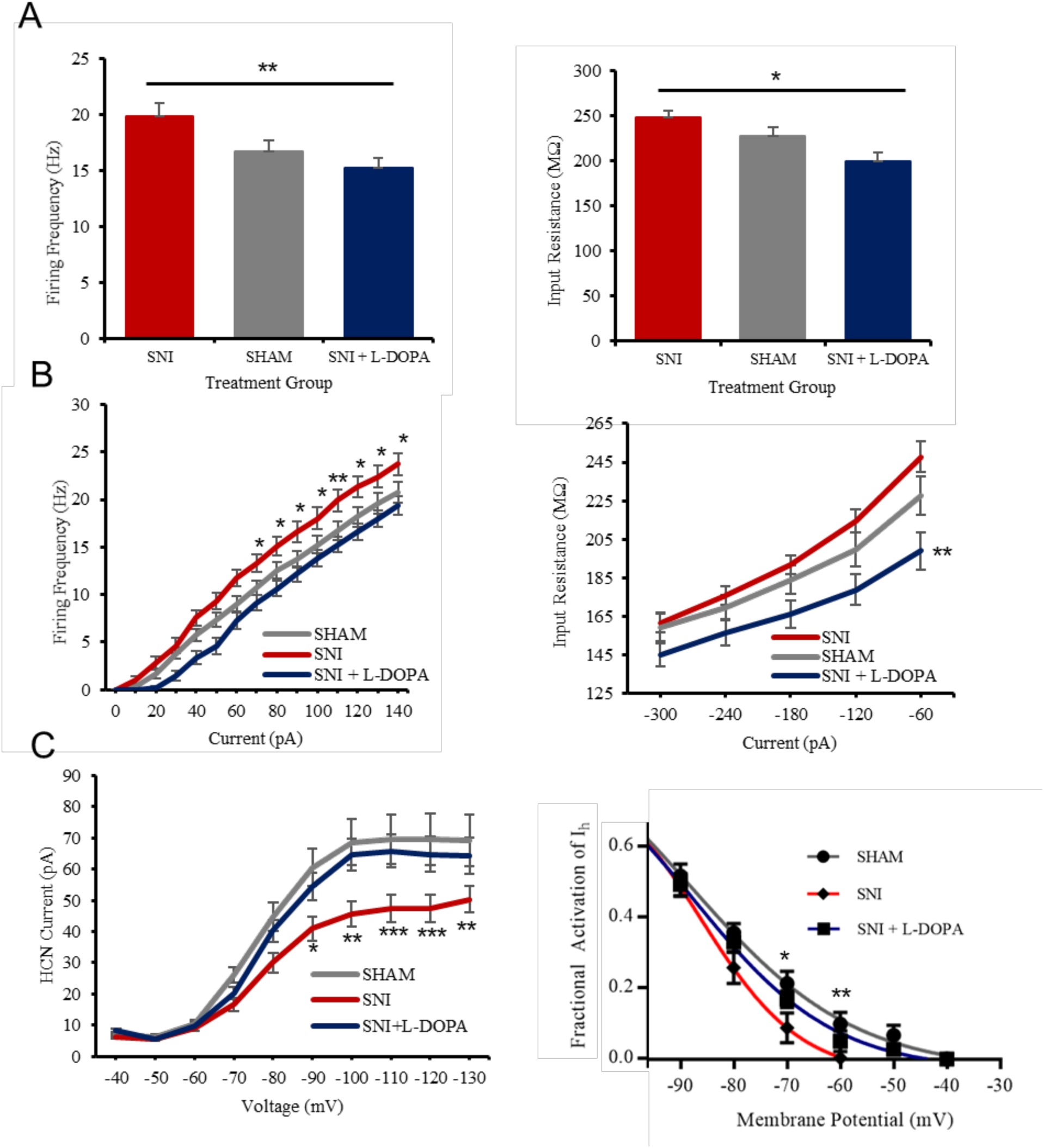
L-DOPA/carbidopa treatment rescues pyramidal excitability and HCN channel function in the ACC of neuropathic mice. A. Pyramidal cells in the ACC of SNI mice under L-DOPA/carbidopa regimen display significantly lower excitability and input resistance than those in control SNI and sham mice. B. Input-output plots documenting L-DOPA/carbidopa-induced decrease in both pyramidal excitability (left) and input resistance (right). C. Rescue of HCN channel activity in L-DOPA/carbidopa-treated SNI mice, with open channel probabilities on level with sham mice at resting membrane potential. Values represented as means ± SEM (n = 13-28): *p < 0.05, **p < 0.01, ***p < 0.001, ****p < 0.0001.

To determine if DA is involved in mediating I_h_, we used voltage-clamp electrophysiology to record HCN currents, in presence of TTX and BaCl_2_ to block voltage-dependent Na and K_ir_ channels. As expected, our recordings indicate that application of 50 µM DA does not cause any significant changes in maximal I_h_ currents measured at −130 mV. However, we noticed a significant increase in I_h_ around physiological membrane potentials (ex. −70 mV) (Figure 4A). A large population (64.2%) of pyramidal neurons in the ACC showed an increase in I_h_ in response to DA (n = 9 of 14). At −70 mV, I_h_ increases from 18.4 ± 2.7 pA to 28.0 ± 3.1 pA following application of 50 µM DA (Figure 4A). This modulation of I_h_ was found reversible as washing-out of DA with aCSF for 10 minutes brings back I_h_ to baseline values (n = 10; paired t-test: t(9) = 3.8, p = 0.0022). A voltage-dependent increase in I_h_ is associated with an increase in fractional activation of HCN channels rather than in the number of HCN channels present at the cell surface (Biel et al, 2009). To check if DA induces an increase in HCN open channel probability, we analyzed HCN tail currents before and after DA application (Figure 4B). We found a significant increase in fractional activation of I_h_ (n = 5; repeated measure ANOVA: F(9, 40) = 64.7, p < 0.0001) (Figure 4B). Consequently, DA induces a rightward depolarizing shift in the half-activation potential of HCN channels, from −89 ± 3 mV to −68 ± 8 mV (Figure 4B) (n = 5; paired t-test: t(8) = 2.5, p=0.0390), indicating that DAergic inhibition is mediated by the modulation of HCN channel activity in pyramidal cells of the ACC.

### DAergic Inhibitory Transduction Remains Functional in Neuropathic Conditions

To determine whether DA-mediated pyramidal inhibition is still effective in ACC in neuropathic conditions, we tested the effects of DA and SKF81297 on pyramidal excitability in mice 4 weeks post-SNI surgery. We observed that 36% and 50% of layer 2/3 ACC pyramidal neurons responded to DA and SKF81297, respectively, values similar to the percentage of responsive cells in naïve animals. SNI mice with tactile and cold allodynia displayed significant inhibition in response to 50 µM DA application, similar to naïve mice (Figure 5A) (n = 4; paired t-test: t(4) = 2.65, 3.00; p = 0.038, 0.029 for SNI and sham respectively). Additionally, 10 µM SKF81297 decreased pyramidal excitability to 76.2 ± 2% of baseline values in both SNI and sham mice. (Figure 5B) (n = 4-6; paired t-test; t(4-5) = 3.33, 15; p = 0.029, 0.0006; for SNI and Sham respectively). At +40 pA stimulation, both 50 µM DA and 10 µM SKF81297 application resulted in full silencing of responsive pyramidal neurons (data not shown). Excitability of pyramidal neurons not exposed to DA or SKF81297 stayed within ∼ 2% of baseline values in both SNI and sham mice (Figure 5B).

### L-DOPA/Carbidopa Administration Rescues Mechanical and Cold Allodynia

Our electrophysiological results, showing that the DA inhibition mechanism is preserved in neuropathic conditions, suggest that DAergic input to the ACC could be diminished in neuropathic conditions. Therefore, we tested the effects of long-term administration of the DA precursor L-DOPA, with carbidopa to prevent its peripheral metabolism, in neuropathic mice. We measured sensory thresholds in neuropathic mice using the von Frey assay and acetone cold test to measure tactile and cold allodynia, two somatosensory hallmarks of chronic neuropathic pain. 2 weeks following SNI surgery, with continuous L-DOPA (75 mg/24 hr/kg) and carbidopa (10 mg/24 hr/kg) intake, we observed treated mice did not display any significant ipsilateral tactile hypersensitivity when compared to the control SNI group (n = 7-11; repeated measure two-way ANOVA with Dunnett’s multiple comparisons: p = 0.0068) (Figure 6A). Similarly, L-DOPA/carbidopa-treated neuropathic mice did not display ipsilateral allodynic responses in the acetone cold test (Figure 6B).

To test whether L-DOPA is effective at reducing hyperalgesia only when provided during a critical time window following SNI surgery, we delayed the start of L-DOPA treatment in one group of mice until 2 weeks following the surgery (SNI + Delayed L-DOPA). Fourteen days following surgery, we tested these mice for tactile allodynia and did not show any significant difference with the control SNI group. After 7 days on L-DOPA, these mice still did not differ significantly from the SNI group, although an upward trend was noticed. Finally, following two weeks on L-DOPA and 4 weeks post-SNI surgery, these mice were statistically identical to the sham, L-DOPA-treated sham and L-DOPA-treated SNI groups and significantly more hypoalgesic than the control SNI group (n = 8; repeated measure two-way ANOVA with Dunnett’s multiple comparisons: p = 0.0120) (Figure 6A). As a control, we added a group of mice who had undergone a sham surgery and supplemented their diet with L-DOPA/carbidopa. The L-DOPA-treated sham group did not differ significantly from the non-treated sham group (n = 6, 10) (Figure 6A). Also, no tactile or thermal allodynia was detected on the contralateral paw in any of the test groups (Figure 6B). Consistent with established literature, mice supplemented with L-DOPA/carbidopa displayed significant motor side-effects, including hyperactivity, starting 5 weeks following the onset of the treatment (Lundblad et al, 2005). No behavioral experiments were conducted on mice showing motor symptoms related to L-DOPA.

### Daily L-DOPA/Carbidopa Administration Rescues Excitability and HCN Function in Neuropathic ACC

In order to assess how the behavioral characteristics of neuropathic mice translate to physiological properties of pyramidal neurons in the ACC, we analyzed the firing frequencies, input resistance, and HCN properties of layer 2/3 pyramidal neurons of SNI mice and SNI mice supplemented with L-DOPA/carbidopa. Four weeks following SNI surgery, with continuous L-DOPA (75 mg/24 hr/kg) and carbidopa (10 mg/24 hr/kg) intake, we observed in patch clamp recordings that neuropathic mice did not display cortical hyperexcitability (Figure 7A). ACC pyramidal cells of L-DOPA-treated SNI mice had firing frequencies on par with sham mice, significantly lower than SNI mice not supplemented with L-DOPA. At + 100 pA stimulation, pyramidal cells from control SNI mice without L-DOPA fired at 18.0 ± 1.1 Hz whereas pyramidal cells from L-DOPA-treated SNI mice fired at 13.8 ± 0.8 Hz (Figure 7A). This effect of the L-DOPA treatment on excitability is also illustrated by the input-output plot (Figure 7B) (n= 13-27; repeated measure two-way ANOVA with Dunnett’s multiple comparison: at 100 pA p= 0.0274). The impact of the L-DOPA treatment of mice on excitability of pyramidal cells in the ACC is logically reflected in changes in their input resistance (Figure 7A and 7B, from 247.8 ± 7.9 MΩ in SNI to 199.0 ± 9.7 MΩ in SNI mice supplemented with L-DOPA/carbidopa (n = 9, repeated measure two-way ANOVA with Dunnett’s multiple comparison: p = 0.0025).

As HCN channel dysfunction is a major driving force causing cortical hyperexcitability in SNI mice, we also measured I_h_ and the fractional activation of HCN channels in neuropathic conditions with or without treatment with L-DOPA. As we have previously reported (Cordeiro Matos et al, 2015), we measured a significant shunting of pyramidal I_h_ in SNI mice, whereas pyramidal cells of SNI mice supplemented with L-DOPA/carbidopa did not display this downward shift, but rather displayed I_h_ values almost identical to those seen in sham mice across all voltages (Figure 7C) (n= 27-34; repeated measure two-way ANOVA with Dunnett’s multiple comparison). This shift in I_h_ is mirrored in the open channel probability of HCN at physiologically relevant membrane potentials. For example, at −70 mV, 16.9 ± 2.3% of HCN channels are open in the SNI + L-DOPA group whereas only 8.8 ± 4.2% of HCN channels are open in the control untreated SNI group (Figure 7C) (n= 25-28; repeated measure two-way ANOVA with Dunnett’s multiple comparison: p= 0.07).

### Intracortical Injection of D1-like Agonist SKF81297 Produces Conditioned Place Preference in Neuropathic Rats

To assess the unique role of ACC in the effects of supplementation with L-DOPA/carbidopa on pain control, the behavioral effects following microinjection of SKF-81297, a dopamine D1 agonist, in the ACC were investigated in adult male Sprague Dawley rats with SNI-induced chronic neuropathic pain and in sham controls. Bilateral microinjection of SKF-81297 in the ACC (0.5 μg/0.5 μl) had no effect on tactile responses in either SNI or sham rats (Figure 8A). We evaluated the effects of SKF81297 in a conditioned place preference (CPP) learning paradigm that captures the positive perception of relief of ongoing pain (King et al, 2009). In this model, only neuropathic rats spent significantly more time in the chamber that had previously been paired with intracortical injection of SKF81297 (Figure 8B and 8C) (repeated measure ANOVA with Sidak’s multiple comparisons test: p < 0.001). Intracortical injection of SKF81297 in sham-operated animals did not produce a chamber preference (Figure 8B). As illustrated in Figure 8C, there is a significant difference of CPP between sham and SNI animals before and after conditioning (n = 15 – 17; unpaired t-test: t(30) = 3.1, p = 0.0047).

**Figure 8.**
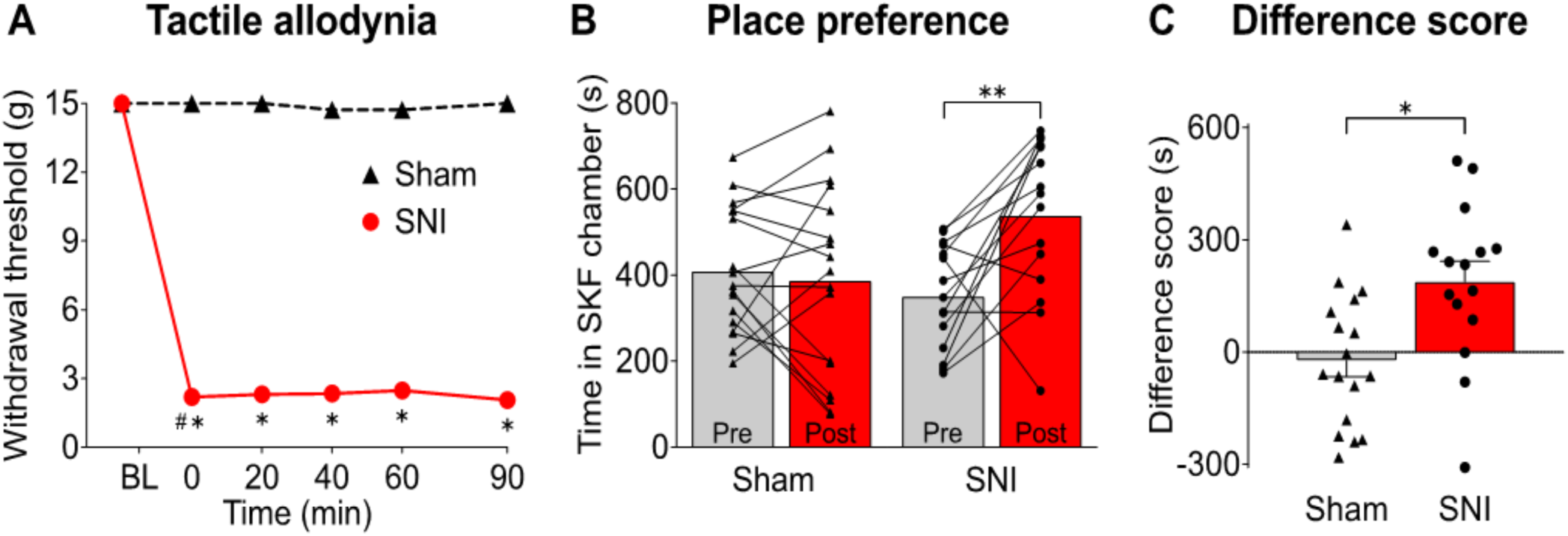
Conditioned place preference is elicited by injection of the selective D1-like agonist SKF81297 in neuropathic ACC. A. Bilateral microinjection of SKF81297 in the ACC (0.5 µg/0.5 µl) had no effect on tactile responses in either SNI or sham rats. Paw withdrawal thresholds were significantly lower in SNI rats compared to shams at all time points during the 90 min testing post SKF81297 treatment. B. In the CPP paradigm, SNI rats spent significantly more time in the SKF81297 paired chamber after drug conditioning (Post) compared to the time in the same chamber before conditioning (Pre). Sham animals did not show significant changes in chamber preference. C. CPP difference scores, calculated as the difference between post-conditioning and pre-conditioning time spent in the SKF81297-paired chamber, demonstrate a significant difference between sham and SNI groups. Values represented as means ± SEM, symbols show the values for individual animals (n = 12-17): *p < 0.0001, **p < 0.01.

## Discussion

Understanding how chronic pain impacts cortical circuitry is key to the development of improved effective treatments for patients. Modulation of ACC neuronal activity by DA is of particular interest in this search given the key role of ACC in the processing of affective components of pain perception, the presence of a dense mesocortical DAergic projection, and the expression of DA receptors across all cortical layers (Clarkson et al, 2017). Our findings indicate that DA is inhibitory to a large subpopulation of pyramidal neurons in ACC both in physiological and neuropathic pain states. DAergic inhibition of ACC neurons raised the rheobase by approximately 30 pA, a difference that could determine whether a neuron fires or not following pain stimuli-evoked postsynaptic potentials in the ACC *in vivo* (Sikes et al, 1992). We used selective D1-like and D2-like ligands to determine that only D1-like receptors are involved in this inhibition (Clarkson et al, 2017). The inhibition induced by the selective D1-like agonist SKF81297 mimicked the inhibition induced by DA while the selective D2-like agonist quinpirole evoked an increase in pyramidal excitability. Additionally, we show that the D1-like antagonist SCH23390 antagonizes DA’s inhibitory effects. D1-like receptors are coupled to Gs proteins and activate the cAMP/PKA pathway. Although it is arguable that these DA-evoked excitatory changes are not present in vivo due to the upregulation of AMPA receptors via PKA phosphorylation, DA has been shown to decrease evoked-postsynaptic potentials (EPSPs) in the ACC (Darvish-Ghane et al, 2016; Esteban et al, 2003; Sun et al, 2005).

The ACC plays a pivotal role in working memory and patients suffering from chronic pain commonly report impairments in working memory performance (Schnurr et al, 1995; Bradley et al, 1989). The property of persistent firing observed in pyramidal cells of the ACC provides a cellular substrate for working memory (Kaminski et al, 2017). We used a glutamatergic model of persistent firing evoked by the mGluR agonist DHPG to test the effects of both DA and SKF81297 on the frequency of action potentials during the persistent depolarization (plateau potential). Our results indicate that SKF81297 and DA consistently decrease the duration of plateau potential and the frequency of persistent firing in the ACC. Thus, the loss of working memory performance measured in chronic pain disorders and the overall effect dopamine and D1-like antagonists on working memory could be related to this strong DAergic modulation of persistent firing in the ACC (Cai et al, 1997; Bradley et al, 1989; Kamiński et al, 2017; Panegyres 2004; Sawaguchi et al, 1991).

The dependence of DAergic inhibition on G_s_-coupled D1-like receptors and not on G_i_-coupled D2-like receptors led us to conclude that cAMP plays a pivotal role in controlling the excitability of layer 2/3 ACC pyramidal neurons. Our group and others have shown that cAMP-gated HCN channel dysfunction plays a major role in promoting cortical hyperexcitability in neuropathic conditions (Cordeiro Matos et al, 2015; Santello et al, 2017). We conclude that the firing activity of pyramidal cells in the ACC is controlled by a balance of monoaminergic inputs through their G_s_ and G_i_-coupled receptors with opposite effects on HCN channel function: activation of G_i_-coupled α_2_ adrenergic receptors shuts down HCN channels while activation of G_s_-coupled 5-HT7 and D1-like receptors increase the opening of HCN channels (Santello et al, 2017; Zhang et al, 2013).

D1 receptors are co-localized with HCN channels on dendritic spines of pyramidal neurons and we confirmed that DA modulates HCN channels through D1-like receptors (Paspalas et al, 2013). Interestingly, our results show D1-like receptors induce an increase in resting membrane potential which is prevented when HCN is blocked with ZD7288; this is likely due to inward cationic currents mediated by HCN channels. This could paradoxically increase the excitability but it is unlikely extracellular DA will be in such high concentrations for long periods of time *in vivo*. Opening of HCN channels induces a decrease in input resistance stemming from a reduction in charge insulation in dendritic spines (Surges et al, 2004). Therefore, the decrease in input resistance induced by D1-like receptor activation is likely overriding this slow depolarization. Although DA does not increase the maximal amplitude of I_h_, we noticed a significant increase in HCN-mediated currents at physiological membrane potentials. Accordingly, the half-activation potential, the membrane potential at which half of HCN channels are open, decreases drastically in the presence of DA. This suggests a change in internal cAMP levels rather than a change in the number of HCN channels at the cell surface. These results are also reflected in the fractional activation of I_h_ at physiological membrane potentials; at −60 mV, DA approximately triples I_h_ fractional activation.

HCN dysfunction appears to be driving cortical hyperexcitability in neuropathic conditions and our findings indicate that D1-like receptors have a strong hold over HCN function. Therefore, defective DAergic inputs to the ACC, creating an offset in the metabotropic control of input resistance in pyramidal cells, could contribute to the neuropathic pain phenotype.

We provide evidence that the signaling mechanisms underlying D1R-mediated inhibition are still functional in the ACC of neuropathic animals, strongly suggesting hypoactive dopaminergic inputs. Interestingly, pathologies with hypoactive supraspinal DAergic pathways, such as Parkinson’s Disease (PD) and major depression, show a high prevalence of non-myogenic chronic pain (Blanchet et al, 2018; Defazio et al, 2008; Djaldetti et al, 2004; Ford 2010). The VTA and the SNc, the main sources of mesocortical DA, display altered firing frequencies and cell death in depression and PD, respectively (Alexander 2004; Liu et al, 2018; McGeer et al, 1988). Furthermore, there is evidence linking a GABA-mediated reduction in dopaminergic activity in the VTA of rodents afflicted with chronic neuropathic pain (Huang et al, 2019; Ko et al, 2018). A reduction in cortical DA could be causing the high comorbidity of chronic pain in PD patients as fMRI studies display pathological hyperactivity in the ACC (Brefel-Courbon et al, 2005). Administration of L-DOPA, a DA precursor that increases DA release in the cortex, and carbidopa, a peripheral DOPA decarboxylase inhibitor, alleviates symptoms of chronic pain and normalizes ACC hyperactivity in PD patients (Antinori et al, 2018; Brefel-Courbon et al, 2005; Schestasky et al, 2007). There is also increasing evidence linking other hypodopaminergic disorders, such as clinical depression, to chronic pain (Fishbain et al, 1997; Taylor et al, 2016).

To test this idea of an hypodopaminergic state in chronic pain conditions, we supplemented chronic neuropathic mice with systemic L-DOPA and carbidopa, a treatment that was successfully used to treat pain in PD patients (Brefel-Courbon et al, 2005). Two weeks following the start of L-DOPA/carbidopa administration, mice that had undergone SNI surgery did not show the typical hypersensitivity of animals in chronic pain as expected tactile and cold allodynia did not differ from sham-operated controls. Additionally, delaying the L-DOPA treatment until 2 weeks post SNI showed reversal of established tactile hypersensitivity. These data suggest that the establishment of chronic pain is associated with a hypodopaminergic state that persists for an extended period. Administration of a L-DOPA both prevented and reversed established behavioral consequences of neuropathic pain suggesting important therapeutic opportunities. Although carbidopa was present to limit the peripheral effects of L-DOPA, we confirmed these behavioral changes are consistent with a reduction in hyperexcitability and input resistance of pyramidal cells in the ACC to levels on par with sham-operated mice and are not peripherally driven. We also measured a dramatic increase in I_h_ at relevant membrane potentials in L-DOPA-treated SNI mice, particularly at resting potentials where there are no HCN channels open in SNI conditions.

Consistent with the demonstration that altering tonic DA levels in the brain over long periods of time has a dramatic normalizing effect on the electrophysiological properties of pyramidal neurons and on the behavioral phenotype of chronic pain animals, we also observed a prominent CPP effect of the acute activation of D1-like receptors with SKF81297 in the ACC of neuropathic pain in a second rodent species. As the CPP paradigm reflects learning resulting from reward associated with pain relief, our findings indicate that DA has analgesic properties in chronic pain conditions (King et al, 2009; Porreca et al, 2017). We did not notice SKF81297-paired chamber preference in sham animals, in agreement with our results on the sham L-DOPA-treated group which did not show any further hypoalgesia than the sham group. This seems to hint at a ceiling effect: a lack of DAergic signaling in chronic pain conditions removes the brakes from the excitability machinery driven by HCN channels, yet providing more DA in non-neuropathic conditions does not have an effect on pyramidal cell activity (Ko et al, 2018). We noticed that intracortical injection of SKF81297 did not have a significant effect on tactile responses in either sham or SNI rats. These results are consistent with previous studies reporting no changes in sensory thresholds with direct optogenetic inhibition of ACC pyramidal neurons (Sellmeijer et al, 2018). Consequently, this suggests that the alleviation of tactile allodynia induced by L-DOPA is not mediated by the ACC. The nucleus accumbens (NAc), a key region of the mesolimbic pathway linked to the somatosensory component of pain, has been shown to be hypodopaminergic in chronic pain due to lower VTA activity (Chang et al, 2014; Ren et al, 2015). Decreasing the excitability of NAc D2+ projection neurons via either chemogenetics or DAergic agonists suppresses the tactile allodynia present in neuropathic pain (Pelissier et al, 2006; Ren et al, 2015; Sarkis et al, 2011). Systemic administration of L-DOPA may therefore normalize DA input to the NAc and the ACC, thereby modulating both the sensory and affective aspects of neuropathic pain.

Although it remains unclear whether hyperexcitability in the ACC is a cause or effect of chronic pain state, increasing evidence suggests inhibiting pyramidal cells of the ACC induces analgesia in chronic pain conditions (Kang et al, 2015; Kuner et al, 2017; Sellmeijer et al, 2018). Our findings strongly suggest that DA is a major inhibitory neuromodulator of pyramidal activity in the ACC in normal as well as in pathological states. We propose that dysregulation of this DAergic inhibition contributes to the expression of typical affective and somatosensory features of chronic pain and that normalization with DA agonists may provide significant therapeutic benefit.

## Materials and Methods

### Subjects and SNI Model

Six-week old male C57BL/6 mice (Charles River Laboratories, QC, CA) were housed in the Montreal Neurological Institute (MNI) Animal Care Facility (ACF) and all procedures are following the Canadian Council on Animal Care guidelines. Male Sprague-Dawley rats, 250-300g (Harlan Laboratories, Indianapolis, IN), were used for CPP experiments. Spared nerve injury (SNI) surgery to induce chronic neuropathic pain was performed on mice and rats 14 days prior to CPP and von Frey testing. Isoflurane (2%) and ketamine/xylazine (80/12 mg/kg i.p.) were used to anesthetize mice and rats, respectively.

### Electrophysiological Recordings

Animals were anesthetized with an Avertin solution (2.5 g tribromoethanol in 5 mL amylene hydrate diluted in 100 mL _dd_H_2_O) (Sigma-Aldrich, St. Louis, MI). Then animals were transcardically perfused with a 4°C choline-chloride based cutting solution oxygenated with carbogen (O_2_ 95%, CO_2_ 5%) (Praxair). Brains were extracted and sliced into 300 µm sections using a vibratome (Leica VT1000). Slices were allowed to rest at room temperature for 1 hour in oxygenated (see above) artificial cerebrospinal fluid (aCSF) containing 124 mM NaCl, 2 mM KCl, 26 mM NaHCO_3_, 1.8 mM MgSO_4_, 1.25 mM NaH_2_PO_4_, 10 mM Glucose, 1.6 mM CaCl_2_, pH 7.4. Slices were submerged at 30-32°C on the stage of a Zeiss Axioskop microscope continuously perfused with oxygenated (see above) aCSF containing 1.8 mM Kynurenic Acid and 100 µM picrotoxin at a rate of 1 mL/minute. A near-infrared CCD camera coupled to a x63 water immersion objective was used during patch-clamp recordings (Sony XC-75). Cells were patched with BF150-75-10 glass pipettes (∼ 6 MΩ) pulled with a Flaming Brown Micropipette Puller (Model P-97, Sutter Instruments, US). Pipettes were mounted on a MP-225 micromanipulator and filled with an intracellular solution containing 120mM K-gluconate, 10 mM HEPES, 0.2 mM EGTA, 20 mM KCl, 2 mM MgCl_2_, 7 mM diTrisPhosphate-Creatine, 4 mM Na_2_ATP, and 0.3 mM NaGTP (Sutter Instruments, Novato, CA). An Axopatch 200B amplifier and Digidata 1322A interface digitizer were used for data acquisition (Molecular Devices, San Jose, CA). Signals were low-pass filtered at 10 kHz for current-clamp recordings and at 2 kHz for voltage-clamp recordings, digitized at 20 kHz.

Neurons were patched in areas corresponding to layer II/III dACC according to stereotaxic coordinates (Paxinos et al, 2008). Pyramidal cells were identified based on firing frequency, input resistance (MΩ), and spike adaptation to a 1-second 100 pA cuurent injection pulse. Cells with resting membrane potentials (RMP) above −50 mV or below −80 mV were excluded. Series resistance was compensated (≤ 35 MΩ).

Excitability was evaluated by bringing cells to −60 mV RMP and counting action potentials evoked by a 17-step protocol (current input ranging from −20 pA to +140 pA, 10 pA steps). Input-output was calculated based on action potentials evoked at each step. Input resistance (MΩ) was assessed by injecting cells with a 6-step protocol (current input ranging from −300 pA to 0 pA, 60 pA steps). Input resistance was averaged across steps and evaluated across treatments.

HCN currents (I_h_) were measured in voltage clamp mode in the additional presence of 1 µM TTX and 200 µM BaCl_2_ to block voltage-gated sodium channels and inward rectifying potassium channels (K_ir_), respectively. Cells were held at a holding potential of −40 mV followed by a 2.5 second step ranging from −40 mV to −130 mV holding voltage (10 step protocol, 10 mV steps). Following each respective step, cells were held at −130mV for 1 second to record tail currents. Current amplitudes of HCN channels were measured by calculating the difference between the current at the start of each step (during the peak) and at the end of each step during the plateau phase. Physiological I_h_ was measured at −70 mV and max I_h_ was determined at −130 mV holding voltage. Fractional activation of I_h_ was calculated by measuring the amplitude of the tail current after each step and normalized to the max tail current (see equation below). A Boltzmann function was fitted using GraphPad Prism 7 (GraphPad Software).

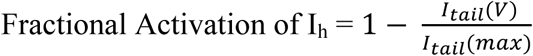

I_h_ amplitude was assessed by obtaining baseline values 5 minutes after patching and comparing them with values obtained following 10-minute treatment with 50 µM DA. Resting membrane potential (mV), temperature, series resistance (MΩ), and whole-cell capacitance were measured before each time point.

### Intracranial ACC Cannulation

Stereotaxic cannulation surgeries were performed in animals anesthetized with a ketamine/xylazine combination (80/12 mg/kg, i.p.; Western Medical Supply/Sigma, Arcadia, CA). Bilateral cannulation of the ACC was performed as previously described (Johansen et al, 2004; Navratilova et al, 2015). A pair of 26-guage stainless steel guide cannulas cut 4 mm below the pedestal (Plastics One Inc., Roanoke, VA) were directed toward the following ACC injection site: AP: +2.6 mm from bregma; ML: ± 0.6 mm; DV: −1.8 mm from skull). Guide cannulas were cemented in place and secured to the skull by small stainless-steel machine screws. Stainless steel dummy cannulas were inserted into each guide to keep the guide free of debris. Rats then received a subcutaneous gentamycin (1 mg/ml) injection, were housed individually, and were allowed to recover for 7-10 days. Cannula locations were verified after the experiment; 9 rats were eliminated due to incorrect placement.

### L-DOPA/Carbidopa Dosage and Administration

Mice were placed on a strict L-DOPA/Carbidopa regimen immediately following SNI surgery. Light-shielded L-DOPA and carbidopa were dissolved in their water supply and administered continually throughout the duration of the study (Blunt et al, 1993). Water intake per mouse was measured 3 times weekly and L-DOPA/carbidopa concentrations were adjusted so as each mouse received 75 mg/24 hr/kg L-DOPA and 10 mg/24 hr/kg carbidopa. In the case of delayed L-DOPA treatment group, administration of L-DOPA/carbidopa started after the first day of behavioral testing instead of immediately following SNI surgery.

### Behavioral Tests

#### Von Frey

Animals were placed in suspended chambers with wire mesh floors for 30 minutes for habituation prior to testing. Mouse withdrawal thresholds were determined by stimulating the lateral planar surface of both the ipsi- and contralateral paws using an automatic von Frey apparatus (Ugo Basile, Italy). Rat withdrawal thresholds were calculated using a series of calibrated von Frey filaments (Stoelting, Wood Dale, IL). Logarithmically spaced increments ranging from 0.41 to 15 g (4–150 N) were applied perpendicular to the plantar surface of the ipsilateral hind paw until the filament buckled. Withdrawal threshold was determined by sequentially increasing and decreasing the stimulus strength (“up and down” method), analyzed using a Dixon nonparametric test, and expressed as the mean withdrawal threshold (Dixon 1980).

#### Acetone cold test

Cold allodynia was measured by applying a drop of acetone to the lateral planar surface of the paw and measuring acute behavioral responses of the mice (0 = no response, 1 = hind paw flinching and stamping, 2 = hind paw licking, 3 = exaggerated response and/or vocalization).

#### Conditioned place preference (CPP)

A single trial conditioning protocol was used for CPP as previously described (King et al, 2009; Navratilova et al, 2013). On preconditioning day, rats were placed into the CPP boxes with free access to all three chambers. Anymaze software was used to determine the time spent in each chamber across 15 minutes. Animals spending more than 80% (720 s) or less than 20% (180 s) of the total time in either chamber were eliminated from further testing (8 rats). On conditioning day, rats with ACC cannulas received a saline injection into the ACC and were placed into a conditioning chamber for 30 min. Four hours later, rats received SKF81297 (0.5 µg/0.5 µl) into the ACC and were placed into the opposite chamber for 30 min. On test day, rats were placed drug free into the middle CPP chamber and were allowed to explore all chambers for 15 minutes; the time in chambers was assessed using the Anymaze software to determine chamber preference. Difference scores were calculated as test time minus preconditioning time spent in the SKF81297-paired chamber.

### Drugs and Solutions

Reagents used for aCSF, cutting solution, internal pipette solution, DA, carbidopa, and xylazine were purchased from Sigma-Aldrich (St. Louis, Missouri). TTX, DHPG, SKF81297, ZD7288, quinpirole, and SCH23390 were obtained from Tocris (Bristol, UK). Levodopa was purchased from Santa Cruz Biotechnology (Dallas, Texas). Ketamine was obtained from Western Medical Supply (Arcadia, CA). All drugs were diluted and aliquoted into single-use samples stored at – 20°C.

### Statistical Analysis

All data were analyzed using Clampfit 10 and statistical analysis was performed using GraphPad Prism 7. Values are represented as either raw or normalized means ± SEM with significance threshold set at p < 0.05. One-way ANOVA with Dunnett’s multiple comparison was used to compare multiple independent groups (aCSF vs 10 µM DA vs 50 µM DA). Unpaired t-tests were used to compare two independent groups whereas the paired t-test was used to compare two related groups (baseline vs treatment). Two-way ANOVA with Bonferroni multiple comparison was used to analyze significance between independent samples under multiple conditions. Significance for fractional activation of HCN channels was tested using repeated measure ANOVA. CPP chamber preference was analyzed using the Anymaze software (Wood Dale, IL).

## Acknowledgements

This work was supported by the Canadian Institutes for Health Research (PJT-1453098, PS) and by NIH (R01 DA 041809, FP). KL holds a studentship from the Louise and Alan Edwards Foundation for Pain Research. We thank our McGill colleagues Stephen Glasgow, Maria Zamfir and Steven Cordeiro Matos for advice and technical support.

